# Sex differences in pain: Spinal cord injury in female and male mice elicits behaviors related to neuropathic pain

**DOI:** 10.1101/2022.10.18.512805

**Authors:** Sydney E. Lee, Emily K. Greenough, Paul Oancea, Ashley R. Scheinfeld, Apsaline M. Douglas, Andrew D. Gaudet

## Abstract

Spinal cord injury (SCI) in humans frequently causes intractable chronic pain. Females are susceptible to worsened pain compared to males, and females may show higher pain prevalence after SCI. Despite this difference in clinical prevalence of SCI pain, few preclinical studies have systematically studied in rodents sex differences in SCI-elicited pain-related behaviors. Here, we leverage data from a large cohort of mice to test whether contusion SCI consistently causes pain symptoms in mice, and to establish whether female (vs. male) mice display heightened hypersensitivity after SCI. Mechanical and heat sensory thresholds were assessed using the von Frey test and Hargreaves test, respectively. In an initial experiment, female mice receiving moderate 60 kDyn SCI or moderate-to-severe 75 kDyn SCI at T9 both exhibited mechanical and heat pain symptoms compared to sham controls. 75 kDyn SCI caused excess motor deficits that confounded defining pain sensitivity at acute times, so the moderate SCI force was used for subsequent experiments. Next, adult female and male C57BL6/J mice received sham surgery or T9 moderate contusion SCI. Comparing female to male mice after SCI, we reveal that mice of both sexes displayed mechanical and heat hypersensitivity compared to sham controls, from acute-to-chronic post-injury times. Females had amplified SCI-elicited hypersensitivity compared to males. Our data suggest that thoracic contusion SCI elicits consistent and persistent pain-associated symptoms, which are more intense in female vs. male mice. These results have important implications for uncovering sex-specific mechanisms and therapeutic targets to ameliorate neuropathic pain after SCI.

## Introduction

Neuropathic pain arises as a secondary complication in 65-80% of patients with spinal cord injury (SCI) (Siddall et al., 1999; Burke et al., 2017). Neuropathic pain is pain caused by damage to the nervous system and presents as a range of pain responses, including allodynia (pain resulting from non-painful stimuli), hyperalgesia, and spontaneous maladaptive pain (Shiao and Lee-Kubli, 2018). Unfortunately, clinical management of neuropathic pain is difficult, and the underlying mechanisms remain poorly understood (Anderson, 2004; Collinger et al., 2013). Accordingly, rodent models of SCI pain have been developed to discover mechanisms and potential therapeutic targets. For instance, midline thoracic contusion SCI in rats or mice causes below-level pain in the hindpaws (Crown et al., 2008; Detloff et al., 2008; Gaudet et al., 2017; Gaudet et al., 2021). Unilateral contusion SCI in the cervical spinal cord of rats or mice also drives neuropathic pain symptoms – both at and below the level of injury (Detloff et al., 2013; Putatunda et al., 2014; Dietz et al., 2022). Thus, effective SCI models that cause chronic pain have been established. Despite this, most SCI studies have not considered sex as a biological variable, so it remains unclear the extent that SCI pain symptoms are experienced by females vs. males.

Women are ∼25% more susceptible to chronic pain than men (Munce and Stewart, 2007), and females have heightened sensitivities to pain in both humans and rodents (Mogil, 2012; Bartley and Fillingim, 2013). Indeed, accumulating evidence suggests that females with SCI are more likely to experience chronic pain – including increased prevalence of self-reported pain (Norrbrink Budh et al., 2003; Andresen et al., 2016) and pain that impairs ability to work (Cardenas et al., 2004). Further, neuropathic pain-associated behavioral outcomes can have sex-specific underlying physiologic mechanisms (Sorge et al., 2015; Rosen et al., 2017), underscoring the importance of studying pain caused by SCI in both sexes. A few studies have assessed sex differences in preclinical models of SCI-elicited pain; however, research using both sexes has shown conflicting results, with some reporting significant sex differences in pain after SCI (Dominguez et al., 2012; Gaudet et al., 2017), and some showing minimal to no apparent sex differences (Walker et al., 2019; McFarlane et al., 2020). Given the widespread use of rodents to unravel neurobiological mechanisms of pain, there is a need to better understand the extent that clinically relevant SCI elicits pain in mice of both sexes.

Here, we aim to establish whether thoracic contusion SCI causes acute-to-chronic neuropathic pain-associated behaviors. In addition, we test the hypothesis that female mice will exhibit exacerbated SCI-elicited pain-related behaviors compared to males. Mice were tested for mechanical allodynia and heat hyperalgesia prior to surgery, and weekly after sham or SCI surgery to 28 days post-operative (dpo). We collate a large dataset that includes mice of both sexes that receive sham or SCI surgery, which illuminates the extent and duration of pain-related symptoms at acute-to-chronic times after SCI. We reveal that SCI elicits neuropathic pain-associated symptoms in mice of both sexes, and that females are predisposed to worsened mechanical and heat pain-related symptoms after SCI.

## Materials and Methods

### Surgery and Animal Care

All housing, surgery, and postoperative care adhered to guidelines set by The University of Texas at Austin Institutional Animal Care and Use Committee. Adult female and male C57BL/6J mice (8-12 weeks old; Jackson Laboratory, Stock 000664) were housed in sex-matched pairs. Food and filtered tap water were provided *ad libitum*, and all animals were maintained on a 12:12 light/dark cycle.

All animals received buprenorphine hydrochloride (0.075 mg/kg; MWI Animal Health, Cat. 060969) analgesic immediately prior to surgery (to reduce acute post-surgery discomfort). Additional daily post-surgery analgesic doses were withheld to limit potential confounding effects on sensitivity-related behavioral tests. A T9 laminectomy was performed on both sham and SCI mice. Next, mice in the SCI group were subjected to contusion SCI using the Infinite Horizon impactor (Precision Systems and Instrumentation) (severity and forces defined below) (Scheff et al., 2003; Gaudet et al., 2016; Gaudet et al., 2021). All surgeries occurred between Zeitgeber time (ZT) 2-10 (lights on at ZT0 and off at ZT12). Incisions were closed using sutures and wound clips. Post-operative care included daily subcutaneous injections of Ringer’s solution (2, 2, 1, 1, 1 mL on the first 5 days post-operative (dpo), respectively; for both sham and SCI mice) to prevent dehydration, and manual voiding of bladders twice daily (shams handled similarly).

### Experiments and mouse numbers

Experiment 1 sought to define an appropriate T9 SCI contusion force (midline injury; 0 s dwell time) for studying neuropathic pain-associated symptoms. Female mice received sham surgery (n=10), 60 kDyn SCI (n=9; force: 62.3 ± 0.5 kDyn; displacement: 597 ± 35 µm), or 75 kDyn SCI (n=10; force: 76.7 ± 0.8 kDyn; displacement: 729 ± 36 µm). One mouse was excluded (died during surgery).

Experiment 2 was designed to assess sex differences in neuropathic pain-related symptoms in female and male mice following moderate T9 SCI (midline injury; 0 s dwell). This experiment combined several independent smaller experiments all on a wildtype C57BL6/J strain. All experiments forming this study used a balanced combination of female/male mice with sham and SCI surgery. All mice were age-matched and experienced the same conditions for housing and testing. Further, experienced researchers performed Hargreaves and von Frey; the researcher performing the tests were consistent throughout each individual study. All Hargreaves testing and studies were completed by one researcher; von Frey testing was completed by two researchers (again, one researcher completed all von Frey testing within an individual study). Mice received sham surgery (female-sham: n=39; male sham: n=48) or 60 kDyn T9 contusion SCI (female-SCI: n=33, force: 62.8 ± 0.5 kDyn; displacement: 498 ± 11 µm) (male-SCI: n=24, force: 63.5 ± 0.7 kDyn; displacement: 494 ± 12 µm). Mice were prospectively excluded from analysis if the Infinite Horizon force/displacement curves suggested a slip or bone hit; if SCI displacement was 100 µm higher or lower than the mean displacement; if mice died during surgery or prior to the experimental endpoint due to ill health/related euthanasia; or if the BMS score was unusually high at 1 dpo (BMS of 3 or greater). 6 female-SCI mice were excluded (3 due to high displacement, 1 due to low displacement, 2 died during surgery), and 8 male-SCI mice were excluded (2 due to high displacement, 1 due to low displacement, 1 died during surgery, 4 died after surgery due to bladder-related complications). All animals received post-operative care as described above.

### Locomotor Testing

Hindlimb motor function was measured using the Basso Mouse Scale (BMS) scores and subscores for locomotor recovery (Basso et al., 2006) prior to surgery and at 1, 3/4, 7, 10, 14, 21, and 28 dpo. Hindlimb recovery was scored on the BMS scale of 0 (complete paralysis) to 9 (normal motor function) by two observers who were blind to treatment conditions. BMS subscores range from 0 to 11. Subscore points are only tallied once mice are plantar stepping frequently or consistently; therefore, mice plantar stepping occasionally or not at all automatically received a BMS subscore of zero.

### Neuropathic pain-related behavioral testing

Testing for neuropathic pain-associated behaviors was performed as previously described (Gaudet et al., 2017; Gaudet et al., 2021). Mice were acclimated to the von Frey and Hargreaves apparatus for two 40-60 min sessions, followed by two pre-surgery tests (average of the two pre-surgery scores are presented here). After surgery, sensory testing occurred weekly. The values for the left and right hindlimb were averaged to calculate a single score for each animal at each timepoint. Testing occurred within 2 hours between ZT2-6. To control for potential confounds of having both sexes tested simultaneously, female and male mice were tested in separate sessions. Both sham and SCI mice from a single sex were tested in the same session. Some SCI mice at 7 dpo (particularly mice with 75 kDyn SCI) did not always plantar place their hindpaw, so we modified our strategy and noted if we had to stimulate the dorsal or lateral aspects of the hindpaw (depending on what part of the paw was presented on the glass or grid). For Hargreaves in the first experiment, ∼60% of mice with 75 kDyn SCI at 7 dpo presented the lateral or dorsal aspect of the paw, whereas ∼5% of mice with 60 kDyn SCI at 7 dpo required stimulating the lateral or dorsal aspect of the hindpaw. All mice from both groups presented the plantar paw surface for testing from 14 dpo until experiment end. Although we were still able to get appropriate withdrawal thresholds from the dorsal/lateral stimulations, the high rate of impaired plantar placement of 75 kDyn SCI mice at 7 dpo led us to use 60 kDyn SCIs for subsequent pain-related studies. Researchers randomized the order of mice in testing chambers to intersperse and remain blind to the treatment group.

#### Von Frey test for mechanical allodynia

Mechanical sensory thresholds were assessed using the simplified up-down (SUDO) method (Bonin et al., 2014) of von Frey testing to minimize stress and time outside of the homecage. Mice were placed in plastic compartments suspended above a wire mesh (Ugo Basile grid platform, Stoelting 57816) and allowed to acclimate for 40-60 min prior to testing. During testing, von Frey filaments (Touch Test Sensory Evaluator, Stoelting 58011) were pressed against the center of the plantar surface of the hindpaw until the filament buckled and held for a maximum of 3 s. A positive response was recorded if the mouse withdrew their hindpaw or flinched in response to the pressure of the filament. In order to minimize animal stress and ensure accurate scoring, von Frey testing occurred the day prior to Hargreaves testing.

#### Hargreaves test for heat hyperalgesia

Heat sensory thresholds (thermal hyperalgesia) were determined using the Hargreaves test (Ugo Basile Thermal Plantar test, Stoelting 55370) (Hargreaves et al., 1988). Mice were placed in a plastic box on top of a glass surface. An infrared heat source (intensity of 25) was applied to the center of the mouse’s hindpaw, and the latency to respond (lifting their paw or flinching) was automatically recorded. Maximum response latency was 30 s; the heat source automatically shut off at this time. Testing on each mouse’s right and left hindpaw was alternated (three tests per time point - total of 6 tests), and each animal was allowed 5 min recovery between each test to minimize sensitivity.

### Statistics

Results were analyzed using two-way or three-way repeated measures ANOVA followed by Holm-Sidak *post-hoc* tests where appropriate. Levene’s test (2017; Team, 2021) was used to assess equality of variances in groups of mice prior to and after SCI. Specific tests used are described in the text for each dataset. Data were graphed using GraphPad Prism.Statistical analysis was performed using SigmaPlot 14.0, and results with *p* < 0.05 were considered significant. All data are plotted as mean ± SEM.

## Results

### Moderate or moderate-to-severe T9 contusion SCI in female mice elicits below-level neuropathic pain-related symptoms

To establish an effective model for studying SCI-elicited neuropathic pain-related behaviors, mice received sham surgery (T9 laminectomy), moderate 60 kDyn T9 contusion SCI, or moderate-to-severe 75 kDyn SCI. For locomotor recovery, SCI at different forces caused graded deficits and recovery of locomotor function (BMS) (BMS: **Fig. 1a**; BMS subscore: **Supplemental Fig. 1**) (surgery x dpo effect: *F*_10,125_=50.88, *p* < 0.001; sham vs. 60 kDyn, *p* < 0.001 from 1-28 dpo; sham vs. 75 kDyn, *p* < 0.001 from 1-28 dpo). 60 kDyn mice showed higher BMS scores vs. 75 kDyn mice from 7 to 28 dpo (*p* < 0.001 at each timepoint); at 28 dpo, average 60 kDyn mouse BMS scores represent frequent steps and mostly/all coordinated stepping, whereas average 75 kDyn mouse scores represent occasional plantar stepping (28 dpo BMS scores: 60 kDyn, 6.0 ± 0.3; 75-kDyn, 3.8 ± 0.3).

**Figure 1.**
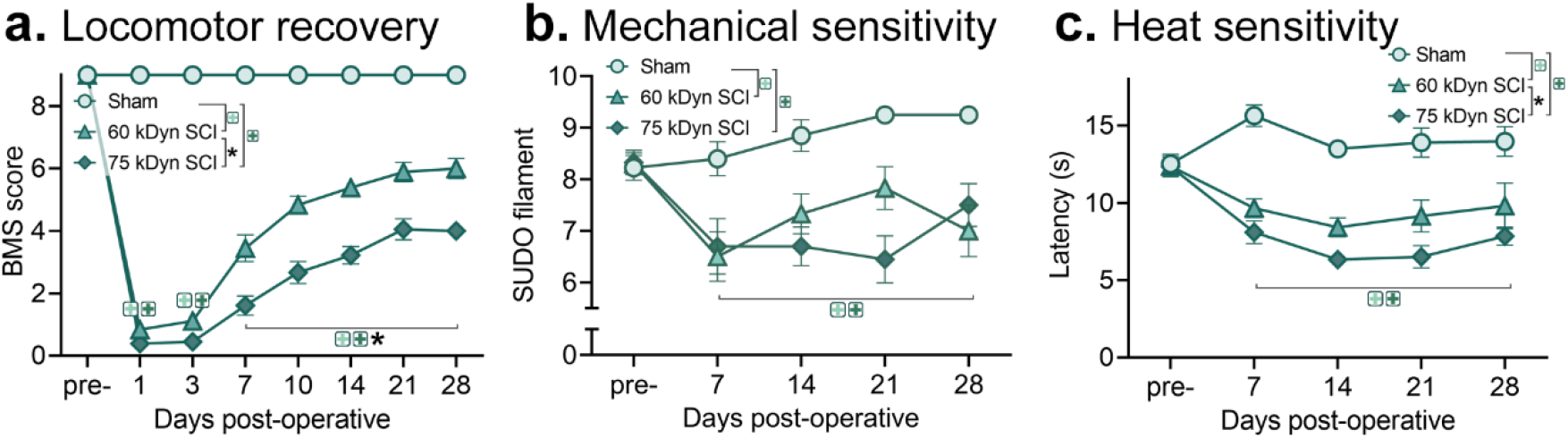
Moderate and moderate-to-severe T9 contusion SCI in female mice cause locomotor deficits and below-level neuropathic pain symptoms. **a**. BMS scores showing SCI-elicited hindlimb locomotor deficits after SCI. As expected, 75 kDyn SCI caused worse deficits than 60 kDyn SCI (from 7-28 dpo) (significant surgery x dpo interaction). **b**. SCI at both severities evoked mechanical hypersensitivity (von Frey test) from 7-28 dpo. **c**. SCI at both severities caused thermal hyperalgesia (Hargreaves test) from 7-28 dpo. Sham: n=10; 60 kDyn SCI: n=9; 75 kDyn SCI: n=10. **+** indicates p<0.05 between sham and SCI mice (color matched to force); * indicates p<0.05 between 60 kDyn and 75 kDyn SCI groups; ANOVA with Holm-Sidak *post-hoc* test.

SCI caused significant and persistent neuropathic pain-related symptoms. Mechanical allodynia was assessed using the von Frey test. Mice in both 60 and 75 kDyn groups exhibited mechanical hypersensitivity at every timepoint between 7 and 28 dpo (vs. sham mice; main effect of surgery: *F*_2,78_=15.54, *p* < 0.001; sham vs. 60 kDyn, *p* < 0.05 from 7-21 dpo and *p* < 0.001 at 28 dpo; sham vs. 75 kDyn, *p* < 0.05 at 7 dpo and *p* < 0.001 from 14-28 dpo) (**Fig. 1b**). There was no significant difference between 60 and 75 kDyn SCI mice in post-SCI mechanical thresholds (*p* > 0.05). Heat hyperalgesia was examined using the Hargreaves test. Both 60 and 75 kDyn SCI mice showed heat hypersensitivity at all times between 7 and 28 dpo (vs. sham mice; main effect of surgery: *F*_2,78_=47.01, *p* < 0.001; sham vs. 60 kDyn, *p* < 0.005 from 7-28 dpo; sham vs. 75 kDyn, *p* < 0.001 from 1-28 dpo) (**Fig. 1c**). There was also a group effect of injury, with 75 kDyn mice showing overall increased sensitivity vs. 60 kDyn mice (Holm-Sidak *post-hoc*: *p* < 0.05). Although the 75 kDyn mice displayed increased heat hypersensitivity – which may suggest this would be a “better” pain model – these mice were more prone to impaired plantar hindpaw placement at 7 and 14 dpo (e.g., see low BMS scores at these times). If the plantar paw surface was not exposed, testers had to stimulate the dorsal or side surfaces of the paw. In these instances, nocifensive responses were not always as clear due to impaired sensorimotor function. Thus, given that 60 kDyn SCI caused comparable pain symptoms to 75 kDyn SCI – but with reduced confounding sensorimotor impairment – the moderate 60 kDyn SCI was chosen for further study of sex differences in SCI pain.

### Moderate contusion SCI causes pain symptoms in mice of both sexes

Next, we used the 60 kDyn T9 contusion SCI validated above to define whether thoracic contusion SCI in mice consistently elicits pain symptoms. In addition, we aimed to assess whether SCI has differential effects on mechanical and heat sensitivity in female vs. male mice. Therefore, a comprehensive analysis of 100+ female and male mice from various studies in the lab were compiled; we reasoned that our large dataset would provide a robust foundation for revealing potential sex differences in SCI-elicited pain symptoms.

First, we evaluated locomotor recovery in female and male mice after SCI (BMS: **Fig. 2a-c**; BMS subscore: **Supplemental Fig. 2**). Mice with sham surgery displayed typical coordinated hindlimb locomotion. Mice with SCI displayed little hindlimb function at 1 dpo (slight hindlimb movement), which recovered to mostly consistent stepping by 28 dpo (BMS: female-SCI vs. female-sham mice – surgery x dpo interaction: *p* < 0.001; male-SCI vs. male-sham mice – surgery x dpo interaction: *p* < 0.001) (both sexes – sham vs. SCI, *p* < 0.001 from 7-28 dpo). Females appeared to have worsened locomotor recovery at 4 dpo and similar long-term recovery compared to males (three-way ANOVA, dpo x surgery x sex interaction, *p* < 0.05). However, when SCI groups were compared, females and males with SCI had no significant difference in locomotor recovery (two-way repeated measures ANOVA: no main effect of sex, *p* = 0.23).

**Figure 2.**
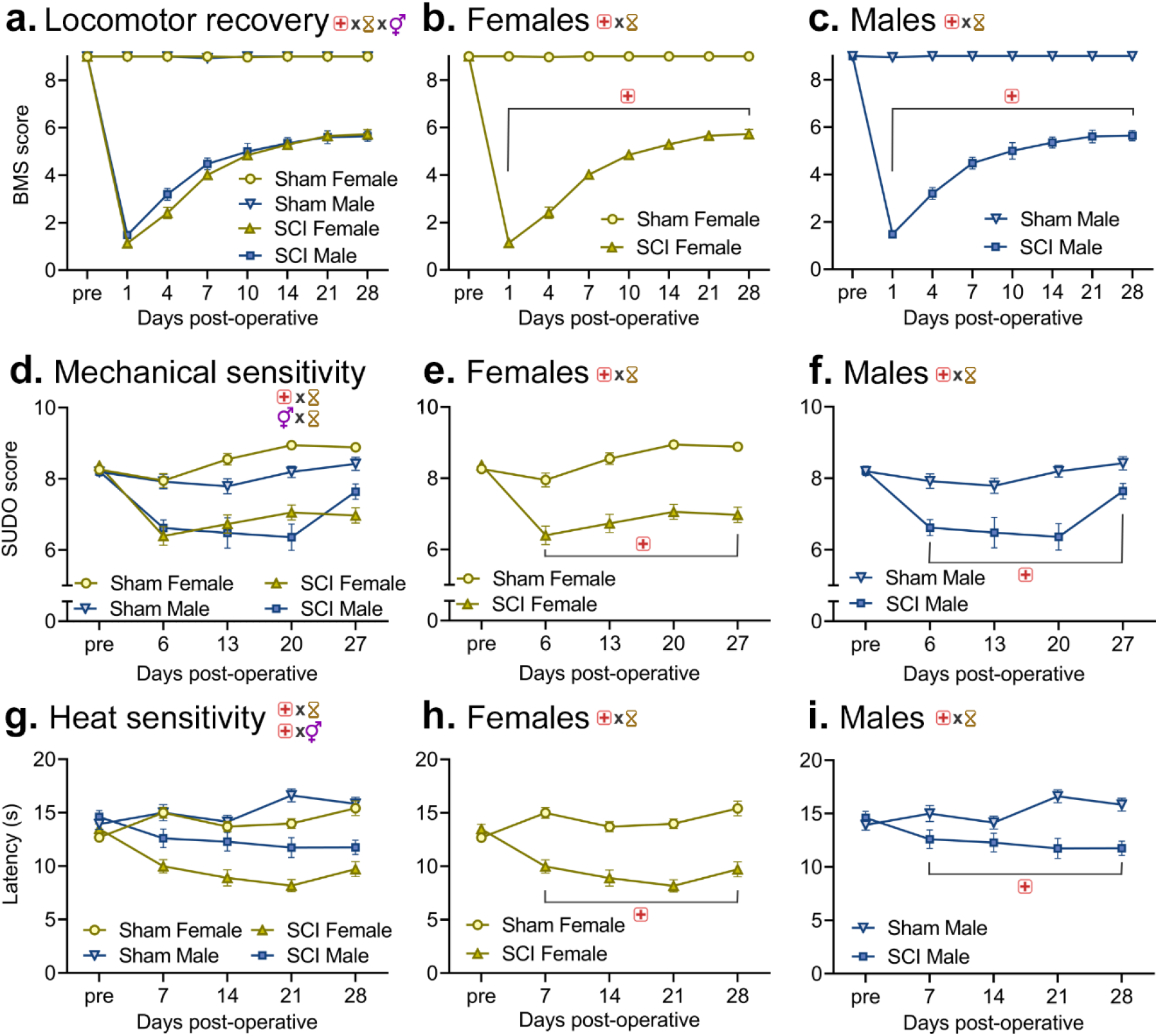
T9 thoracic contusion SCI elicits mechanical and heat hypersensitivity in mice of both sexes, with amplified pain symptoms in females compared to males. **(a-c)** Locomotor recovery after SCI or sham surgery in both sexes (a), females only (b), and males only (c). BMS scores reveal similar recovery curves after SCI between sexes, although females showed slightly delayed recovery at acute times. **(d-f)** Using the von Frey test, female and male mice exhibit SCI-elicited mechanical hypersensitivity compared to sham control mice. **(g-i)** Using the Hargreaves test, female and male mice display heat hypersensitivity after SCI (vs. sham controls). Female-sham: n=39; male sham: n=48; female-SCI: n=33; male-SCI: n=24. + *p* < 0.05 between sham and SCI mice; hourglass symbol indicates *p* < 0.05 across timepoints; gender symbol indicates significant effect of sex; “x” symbol indicates significant interaction between respective groups. Symbols at the top of each panel indicate main effects. ANOVA (a,d,g: three-way; all other panels: two-way repeated measures) with Holm-Sidak *post-hoc* test.

Next, SCI-driven mechanical hypersensitivity was studied using the von Frey test (**Fig. 2d-f**). All groups had similar baseline pre-surgery mechanical thresholds. Mice with sham surgery showed consistently high mechanical thresholds that did not change much over time post-surgery. In contrast, mice with SCI exhibited decreased mechanical thresholds, which represents mechanical hypersensitivity (three-way ANOVA: dpo x surgery interaction, *p* < 0.0001; dpo x sex interaction, *p* < 0.005). Indeed, mechanical hypersensitivity was observed in SCI mice across all post-injury times for females (**Fig. 2e**) and for males (**Fig. 2f**), compared to sham controls (female-SCI vs. female-sham mice – surgery x dpo interaction: *F*_4,220_=15.08, *p* < 0.001; male-SCI vs. male-sham mice – surgery x dpo interaction: *F*_4,216_=5.29, *p* < 0.001) (sham vs. SCI: females, *p* < 0.001 from 7-28 dpo; males, *p* < 0.001 at 7-21 dpo and *p* < 0.05 at 28 dpo). At 28 dpo, females with SCI had persistent, robust mechanical allodynia compared to sham controls, whereas males with SCI had notably reduced magnitude of mechanical hypersensitivity.

To establish whether SCI in mice causes heat hypersensitivity, mice completed the Hargreaves test (**Fig. 2g-i**). SCI elicited significant heat hypersensitivity throughout the post-operative period; heat hypersensitivity was amplified in SCI-females vs. SCI-males (three-way ANOVA: dpo x surgery interaction, *p* < 0.0001; surgery x sex interaction, *p* < 0.05). Females with SCI had reduced latency to withdrawal due to heat at all post-surgery times compared to female-sham mice. Males also had SCI-elicited pain; however, the magnitude of SCI-elicited heat hypersensitivity was reduced compared to females (Female-SCI vs. female-sham mice – surgery x dpo interaction: *F*_4,280_=15.49, *p* < 0.001; male-SCI vs. male-sham mice – surgery x dpo interaction: *F*_4,205_=5.85, *p* < 0.001) (sham vs. SCI: females, *p* < 0.001 from 7-28 dpo; males, *p* < 0.05 at 7-14 dpo and *p* < 0.001 at 21-28 dpo). Therefore, our data suggest that moderate thoracic SCI in mice elicits significant below-level mechanical and heat hypersensitivity.

### Spinal cord injury causes neuropathic pain-associated symptoms that shift over time

The size of our dataset enabled comparing individual patterns between groups and sensitivity thresholds over time pre- and post-SCI (**Fig. 3**). For mechanical sensitivity, pre-surgery thresholds were relatively consistent across mice and had low variability. In contrast, SCI induced higher variability, particularly during the acute-to-subacute times post-injury (Levene’s test for equality of variances; females: *F*_*4,160*_=3.40, *p* < 0.05 pre-surgery vs. 6 and 13 dpo; males: *F*_*4,120*_=6.88, *p* < 0.05 pre-surgery vs. 13, 21, and 27 dpo). For mechanical sensitivity, females with SCI exhibited higher sensitivity (vs. pre-SCI) and a small subset did not show hypersensitivity or were less responsive (**Fig. 3a**). Similarly, males after SCI exhibited mechanical hypersensitivity compared to pre-surgery thresholds, with some SCI males developing intense hypersensitivity at 13-27 dpo (**Fig. 3b**).

**Figure 3.**
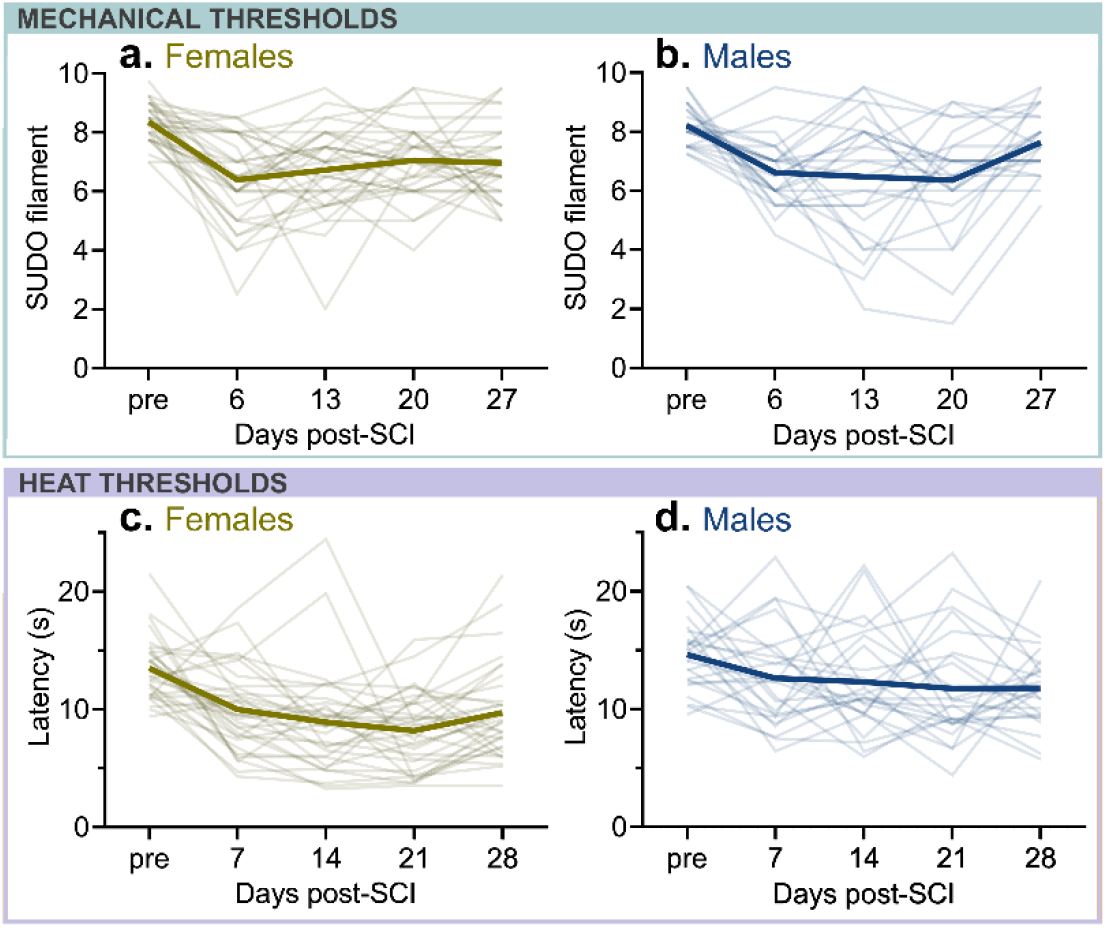
Mechanical and heat sensitivities compared to pre-surgery baseline highlight that SCI induces hypersensitivity in mice of both sexes. For each graph, the light trend lines represent one individual, and the dark trendline is the mean of all individuals. **(a**,**b)** SCI-elicited mechanical hypersensitivity in female (a) and male (b) mice. **(c**,**d)** SCI-elicited heat hypersensitivity in female (c) and male (d) mice. Compared to pre-surgery thresholds, female and male mice with SCI showed increased variability in mechanical thresholds, but not heat thresholds (analyzed using Levene’s test for equality of variances). Female-sham: n=39; male sham: n=48; female-SCI: n=33; male-SCI: n=24.

For heat stimulation, females with SCI again showed hypersensitivity (vs. pre-surgery) with similar variability compared to baseline variability (Levene’s test comparing pre- and post-SCI scores: *p* = 0.52) (**Fig. 3c**). Some SCI females exhibited hyposensation that mostly resolved within 21 dpo. Males with SCI showed heat hypersensitivity, but some male mice showed increased post-SCI variability across days post-SCI (**Fig. 3d**). The paradoxical combination of hypersensitivity vs. loss of sensation due to partial denervation may contribute to variability in individuals’ data after SCI; however, there was no significant overall difference in heat threshold varability after SCI compared to pre-surgery (Levene’s test: *p* = 0.49).

### SCI-driven mechanical pain symptoms are exacerbated in females compared to males

Next, we sought to directly compare pain symptoms in female vs. male mice. To enable comparing females vs. males, “difference scores” were calculated: sensitivity threshold values from SCI mice were subtracted from sham mouse values for each sex (i.e., sham scores minus SCI scores for both mechanical and heat thresholds) (**Fig. 4a**). Accordingly, SCI-elicited hypersensitivity is expressed as an increase on the graph. SCI caused mechanical hypersensitivity in both females and males compared to shams of the same sex (two-way ANOVA: main effect of dpo, *p* > 0.001). Using this metric, there were no significant differences in mechanical hypersensitivity between sexes (*p* = 0.08).

**Figure 4.**
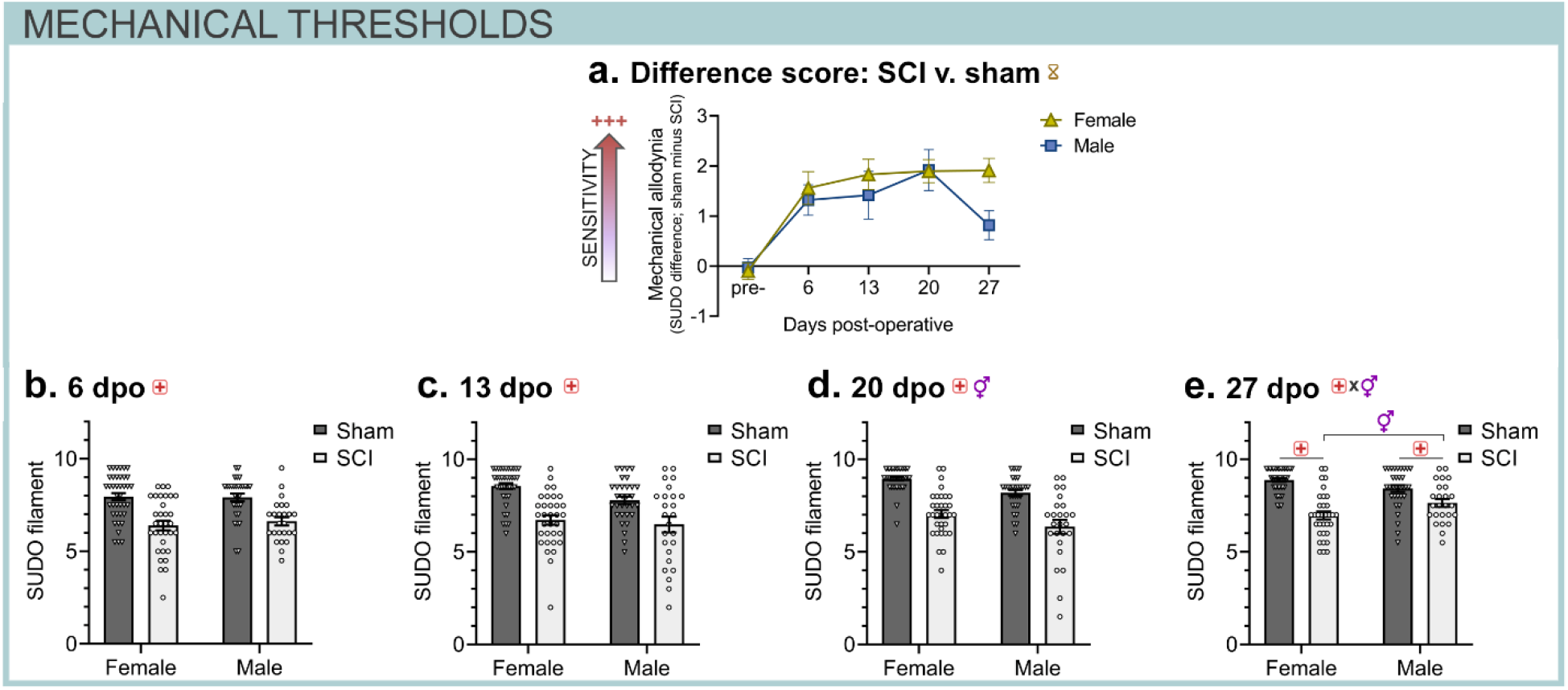
Difference scores and acute-to-chronic thresholds highlight the extent of SCI-elicited mechanical pain symptoms in females and males. **(a)** Difference scores in mechanical sensitivity: subtracting “sham minus SCI” SUDO von Frey scores for each sex shows that both female and male mice exhibit mechanical pain symptoms after SCI (increased pain = upwards on the y-axis). **(b-e)** SCI-driven mechanical allodynia in female and male mice. Individual von Frey scores at 6 dpo (b), 13 dpo (c), 20 dpo (d), and 27 dpo (e). Females exhibit increased SCI-elicited mechanical hypersensitivity compared to males at 20 dpo (main effect of sex) and 27 dpo (after SCI). Female-sham: n=39; male sham: n=48; female-SCI: n=33; male-SCI: n=24. Hourglass symbol indicates *p* < 0.05 across timepoints; gender symbol indicates significant effect of sex; + *p* < 0.05 between sham and SCI mice; “x” symbol indicates significant interaction between respective groups. Two-way repeated measures ANOVA (a) or two-way ANOVAs (b-e) with Holm-Sidak post-hoc test.

Comparing trends in individual mechanical sensitivity threshold values underscores the fact that SCI drives acute-to-chronic mechanical hypersensitivity (**Fig. 4b-e**). SCI elicited increased mechanical sensitivity at all timepoints between 6-20 dpo, to a similar extent between both sexes. At 27 dpo, females and males both displayed SCI-elicited mechanical hypersensitivity compared to sham controls (Two-way ANOVA: sex x surgery interaction: *F*_1,125_=9.89, *p* < 0.005). Further, female-SCI mice had increased mechanical hypersensitivity compared to male-SCI mice (*p* < 0.05).

### SCI-elicited heat hypersensitivity is worsened in females vs. males

SCI also induced heat hypersensitivity (**Fig. 5**). Differences in Hargreaves heat response latencies reveal that SCI drives heat pain symptoms in both female and male mice (compared to sham controls) (**Fig. 5a**) (main effect of sex (*F*_1,305_=6.47, *p* < 0.05) and dpo (*F*_4,305_=14.36, *p* < 0.0001); sex x dpo interaction; *p* = 0.43). Further, notable sex differences in pain behavior were observed. Females had increased heat sensitivity compared to males prior to surgery (*p* < 0.05), which was exacerbated after SCI. Female and male mice with SCI exhibited early, consistent, and persistent heat hyperalgesia compared to sham controls (**Fig. 5b-e**). At 7 dpo, there was no significant effect of sex, although females with SCI appeared modestly more sensitive than males (sex x surgery interaction: *p* = 0.06). Significant sex differences in SCI-induced heat pain symptoms occurred at 14 dpo and 21 dpo. At 14 dpo, females exhibited heightened SCI-elicited heat hypersensitivity compared to males (two-way ANOVA: sex x surgery interaction: *F*_1,123_=6.86, *p* < 0.05; female-SCI vs. male-SCI, *p* < 0.005). At 21 dpo, females had increased heat hypersensitivity compared to males (two-way ANOVA: main effect of sex, *p* < 0.001; main effect of surgery, *p* < 0.001). Overall, our data show that moderate contusion SCI causes below-level pain in female and male mice, and that females have amplified SCI-driven pain symptoms compared to males.

**Figure 5.**
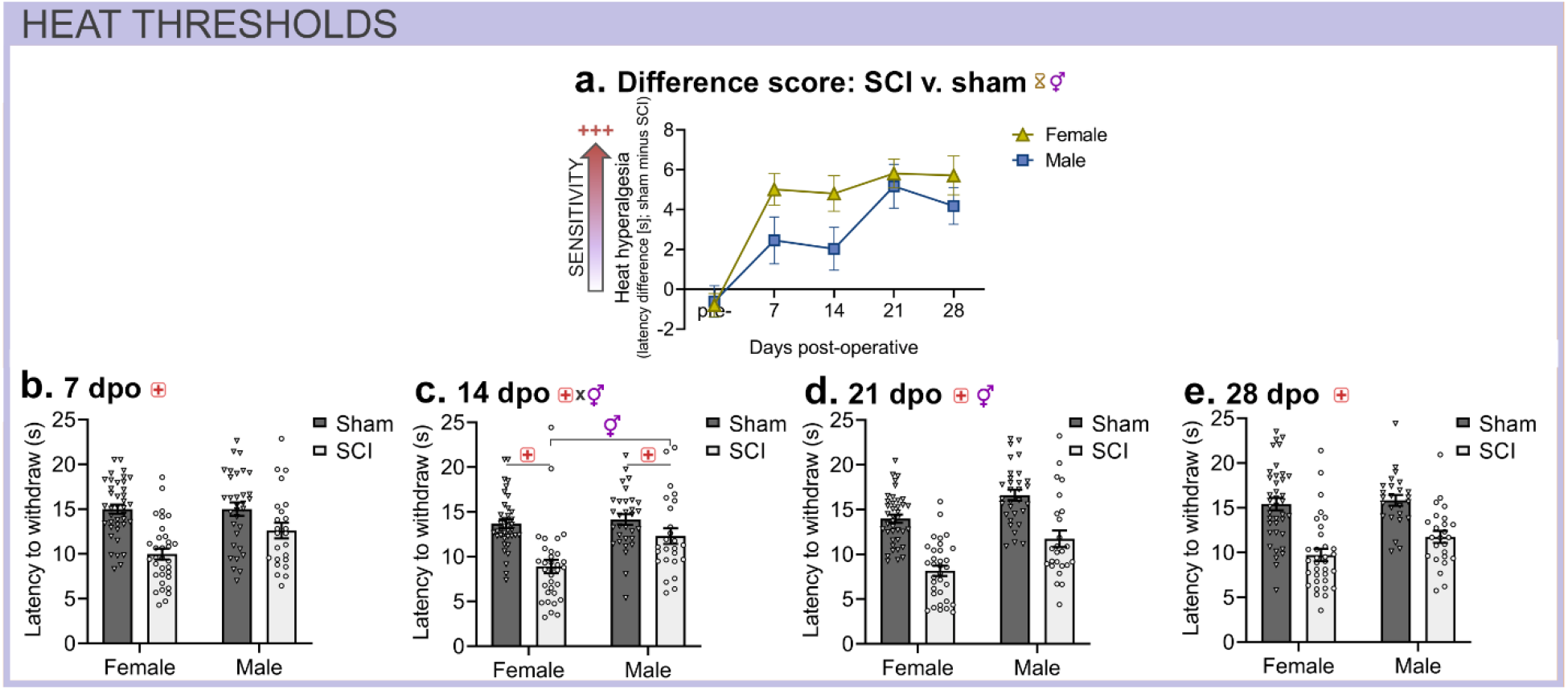
Sex similarities and differences in SCI-evoked heat hypersensitivity in mice: mice of both sexes display heat pain symptoms, and female (vs. male) mice exhibit exacerbated pain symptoms after SCI. **(a)** Difference scores in heat sensitivity: subtracting “sham minus SCI” withdrawal latencies for each sex reveals that both female and male mice exhibit heat pain after SCI (increased pain = upwards on the y-axis). Females had increased SCI-driven pain compared to males. **(b-e)** Individual Hargreaves withdrawal latencies at 7 dpo (b), 14 dpo (c), 21 dpo (d), and 28 dpo (e). Females (vs. males) generally display increased heat sensitivity, and show exacerbated SCI-elicited heat pain symptoms at 14 dpo (after SCI) and at 21 dpo (main effect of sex). Female-sham: n=39; male sham: n=48; female-SCI: n=33; male-SCI: n=24. Hourglass symbol indicates *p* < 0.05 across timepoints; gender symbol indicates significant effect of sex; + *p* < 0.05 between sham and SCI mice; “x” symbol indicates significant interaction between respective groups. Symbols at the top of each panel indicate main effects. Two-way ANOVA with Holm-Sidak post-hoc test.

## Discussion

Here, we leveraged a large dataset to illuminate whether contusion SCI in mice causes neuropathic pain-related behaviors, and whether SCI-elicited pain symptoms present differently in female vs. male mice. Our initial study revealed that two injury forces – a moderate 60 kDyn SCI and a moderate-to-severe 75 kDyn SCI – both caused mechanical and heat hypersensitivity in female mice. The moderate SCI caused fewer paradoxical motor deficits, so was used for subsequent studies. Moderate SCI in female and male mice caused mechanical allodynia and heat hyperalgesia. SCI increased hypersensitivity in pain thresholds within individuals. The extent and duration of SCI-mediated neuropathic pain-related behavior was worsened in female compared to male mice. Overall, our data suggest that SCI in mice causes significant and long-lasting neuropathic pain-associated symptoms; and that female mice are more susceptible to SCI-driven mechanical and heat hypersensitivity.

The purpose of our first experiment was to establish a model of SCI-elicited neuropathic pain-like behaviors. Mice with either 60 or 75 kDyn SCI had mechanical and heat pain symptoms, and the 75 kDyn SCI mice had modestly worsened pain symptoms compared to 60 kDyn SCI mice. This is an important validation study, because it shows that either force can be used experimentally to assess outcomes suggestive of neuropathic pain and its modulation through genetic or therapeutic strategies. We did note that some 75 kDyn SCI mice had excess loss of sensorimotor function at 7 dpo, which masked or confounded establishing sensory thresholds at that timepoint. Further, some 75 kDyn SCI mice at 7 dpo did not always plantar place their hindpaw, so we modified our strategy and noted if we had to heat the dorsal or lateral aspects of the hindpaw (depending on what part of the paw was presented on the glass). Overall, these results corroborate with and extend previous results: we and others showed that thoracic contusion SCI at 60 kDyn (Wu et al., 2016) or 75 kDyn (Gensel et al., 2019; McFarlane et al., 2020; Gaudet et al., 2021) cause pain symptoms in mice, yet no known studies have systematically compared pain that occurs with disparate SCI forces in the same experiment in mice (for force-pain relationships in rats, see (Gaudet et al., 2017). Based on paradoxical sensorimotor impairments in 75 kDyn SCI mice at 7 dpo – an important time for recovery trajectory and epicenter inflammatory dynamics – we used 60 kDyn SCI for subsequent studies herein.

Our results demonstrate that moderate T9 contusion SCI caused mechanical and heat hypersensitivity in mice of both sexes from acute-to-chronic times post-injury. Similarly, others have explored behaviors related to neuropathic pain after SCI in rodents. Detloff et al. (Detloff et al., 2008) revealed that T8 SCI in rats elicited below-level mechanical allodynia similar to at-level allodynia caused peripheral nerve injury. Further, SCI-induced inflammatory activation in the lumbar spinal cord correlated with neuropathic pain-related symptoms. Carlton et al. (Carlton et al., 2009) showed that T10 SCI in rats caused above-level mechanical and heat hypersensitivity, which co-occurred with heightened glial reactivity in the thoracic and cervical spinal cord. In accordance, SCI causes neuropathic pain-associated symptoms in mice, which can be dampened using anti-inflammatory modulators (thoracic SCI: (Wu et al., 2016; McFarlane et al., 2020; Gaudet et al., 2021); cervical SCI: (Brown et al., 2021; Dietz et al., 2022). Others have divided subjects into cohorts that exhibit or do not exhibit pain-related behaviors, then analyzed these cohorts separately (Nesic et al., 2005; Boroujerdi et al., 2011; Chhaya et al., 2019; Dietz et al., 2022). We and others find that this practice is unnecessary using the thoracic contusion injury model in mice; the SCI forces used herein cause significant and reproducible pain-associated symptoms that can be modulated by predictable manipulations (e.g., anti-inflammatory strategies; (Gensel et al., 2019; Gaudet et al., 2021).

We revealed that T9 contusion SCI in mice of both sexes caused pain symptoms, which were magnified in females. Indeed, females had worsened SCI-driven mechanical and heat hypersensitivity compared to males. Others have presented mixed results related to sex differences after SCI. SCI using different models in rats can have divergent sex-specific outcomes (Dominguez et al., 2012; Gaudet et al., 2017). In mice, McFarlane et al. (McFarlane et al., 2020) assessed mechanical allodynia and heat hyperalgesia once every two weeks post-surgery – they found that T9 75 kDyn SCI caused pain symptoms in mice of both sexes to 6 weeks post-SCI, but there were no significant differences between sexes. Interestingly, females showed slightly (but not significantly) increased mechanical hypersensitivity in both von Frey and Hargreaves tests, and 6-9 mice were used per group (McFarlane et al., 2020). This differed from our study in that we used large cohorts of mice, a moderate 60 kDyn injury force, and tested thresholds weekly after SCI. There could also be slight differences across labs due to surgeon technique, the behavioral tester’s experience and technique, and even the gender of the behaviorist (Sorge et al., 2014). Overall, our large cohort, more moderate SCI model, and weekly sensitivity testing has enabled us to deduce that female mice exhibit exacerbated SCI-elicited pain symptoms.

Of note, similar pain outcomes between sexes can have differential underlying cellular and physiologic mechanisms. Peripheral nerve injury in mice causes mechanical allodynia in both sexes. Some studies have revealed sexually divergent cellular drivers of injury-induced pain, suggesting that pain in males is mediated by microglia, whereas pain in females is caused mainly by adaptive immune cells (Sorge et al., 2015; Fernandez-Zafra et al., 2019; Tawfik et al., 2020). Other studies suggest that microglia are required for induction of neuropathic pain-like behaviors in both sexes (Peng et al., 2016), or facilitate pain but on sex-specific post-injury timelines (delayed microglial response in females) (Huck et al., 2022). Neuropathic pain-like behavior can also be elicited by altered excitability of spinal cord circuits via altered dorsal horn neuron excitability or inhibition; dampened descending inhibition; detrimental shifts in excitation-inhibition balance; and altered expression of neuromodulators and immunomodulators (Hulsebosch et al., 2009; Brown et al., 2022). Although sex-specific mechanisms causing SCI pain remain understudied, sex differences in hormones and inflammatory responses after SCI could underpin distinct mechanisms (Stewart et al., 2021; Stewart et al., 2022). Overall, this highlights the importance of incorporating subjects of both sexes throughout all studies (Shansky, 2019; Stewart et al., 2020; Shansky and Murphy, 2021): if no significant sex differences exist then results from both sexes can be combined for analysis, whereas if sex differences become apparent, they can be studied further. Therefore, future studies should include both sexes and will further crystallize our understanding of sex-specific mechanisms that underlie neuropathic pain, which could have crucial implications for development of targeted analgesic therapies.

We limited confounds of performing behavior on both sexes simultaneously by using separate female and male cohorts; however, there are limitations to our approach as well. Testing females and males in separate cohorts required mice being tested on different days and/or times of day. In addition, cohorts of females and males received surgery on different days. Therefore, one limitation is that there could be differences in day/time of testing between sexes that could affect behavioral outcomes. We strived to minimize potential experimental differences by performing female and male studies close in time (e.g., linked cohorts of similar sizes separated by less than a week); by wiping equipment thoroughly between tests with water and then 70% alcohol; by having the same researchers complete testing for mice of both sexes; and by housing and testing female and male mice under the same conditions.

Our study provides an extensive exploration of how SCI in mice of both sexes leads to differences in evoked pain symptoms. Although these acute-to-chronic measures of pain-like behavior aid in identifying relevant mechanisms underlying complex chronic pain conditions, tests that rely on reflexive response do not fully capture the affective components of the pain experience (Kramer et al., 2017). These tests have limits in their utility to identify mechanisms underlying the human neuropathic pain experience. Recently, researchers have been developing relevant pain-like behavioral assays that incorporate affective components of the pain experience, which is a difficult undertaking (Burma et al., 2017).

Promising strategies for measuring affective components of pain include conflict tests, which pair a pain-inducing stimulus against a conflicting stimulus, and studying signs of spontaneous pain behavior (see reviews: (Burma et al., 2017; Kramer et al., 2017). Conflict tests can be used to evaluate differing motivational states through the introduction of approach-avoidance situations. For example, the mechanical-conflict avoidance paradigm is a recently developed conflict test used to assess pain behavior in rats (Chhaya et al., 2019; Gaffney et al., 2022). The apparatus consists of a long rectangular runway connecting a light chamber to a dark chamber. This connecting runway is equipped with small mechanical spikes that produce mechanical sensitivity when crossed. Conditioned place preference has also been used to reveal pain behavior in rats (Yang et al., 2014). Regarding rodent spontaneous pain, the mouse grimace scale was developed to assess mouse pain behavior through differing mouse facial expressions (Langford et al., 2010; Mogil et al., 2020), including after SCI (Wu et al., 2016; Schneider et al., 2017; Heinsinger et al., 2020). Pain-like behavior is assessed using machine learning to identify characteristic orbital tightening, cheek bulge, nose bulge, and whisker position; however, its effectiveness may be limited by animals’ tendency to hide behaviors that signal pain (Burma et al., 2017). Another challenge with assessing SCI-driven pain is that the injury simultaneously affects sensory function (paradoxical loss of sensation and/or pain) and motor function. This phenomenon can confound interpreting results of promising pain-related tests that assume similar motor function across mice/groups. Overall, conflict tests and spontaneous pain measures may complement reflexive pain assays in future studies to better understand the complex mechanisms underlying pain behavior in chronic pain conditions.

## Conclusions and future directions

We employed a commonly used, clinically relevant contusion SCI model to establish whether SCI elicits neuropathic pain-related symptoms in female and/or male mice. We found that both moderate and moderate-to-severe SCI in mice caused neuropathic pain-associated behaviors. Further, our data suggest that midline thoracic SCI in mice causes persistent neuropathic pain-related behaviors in mice of both sexes, and that females are predisposed to worsened sensitivity to mechanical and heat stimuli. Given the high prevalence of pain after SCI in humans, there is a need to develop and validate rodent models that assess SCI-induced neuropathic pain-associated behaviors. Our study provides a crucial foundation for preclinical research related to sex and SCI; future studies should explore whether SCI-elicited pain-related behaviors in females vs. males is due to similar or distinct cellular mechanisms. Future preclinical SCI studies could combine complementary nociceptive tests, validated for SCI here, with newer but more complex strategies that incorporate the affective aspect of the pain experience. Development of clinically relevant models and tests of SCI-elicited pain is critical to identify new therapeutic targets for quenching chronic pain.

## Acknowledgements

We thank the Animal Resources Center (ARC) husbandry staff at the Health Discovery Building for excellent animal care, and Dr. Laura Fonken for helpful critique and feedback on the manuscript. Partial support was provided by University of Texas at Austin start-up funds (ADG), the Wings for Life Foundation (ADG) and Mission Connect, a program of the TIRR Foundation (ADG).

## Notes

*Conflict of interest:* The authors declare no competing financial interests.

### Competing Interest Statement

The authors have declared no competing interest.

### Summary of Updates

Revised to improve clarity of writing and rigor. Added some discussion. Added supplementary data.

