## Supplemental Fig for "Sex differences in pain: Spinal cord injury in female and male mice elicits behaviors related to neuropathic pain"

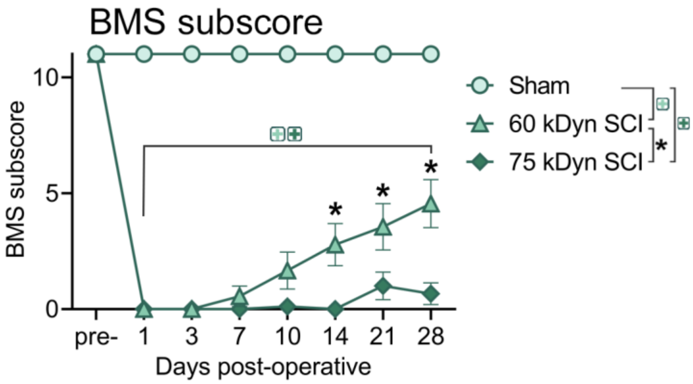


**Supplemental Figure 1.** Moderate and moderate-to-severe T9 contusion SCI in female mice cause graded locomotor deficits as revealed by the BMS subscore. As expected, 75 kDyn SCI caused worse deficits than 60 kDyn SCI (from 14-28 dpo). Sham: n=10; 60 kDyn SCI: n=9; 75 kDyn SCI: n=10. **+** indicates p<0.05 between sham and SCI mice (color matched to force); * indicates p<0.05 between 60 kDyn and 75 kDyn SCI groups; ANOVA with Holm-Sidak *post-hoc* test.

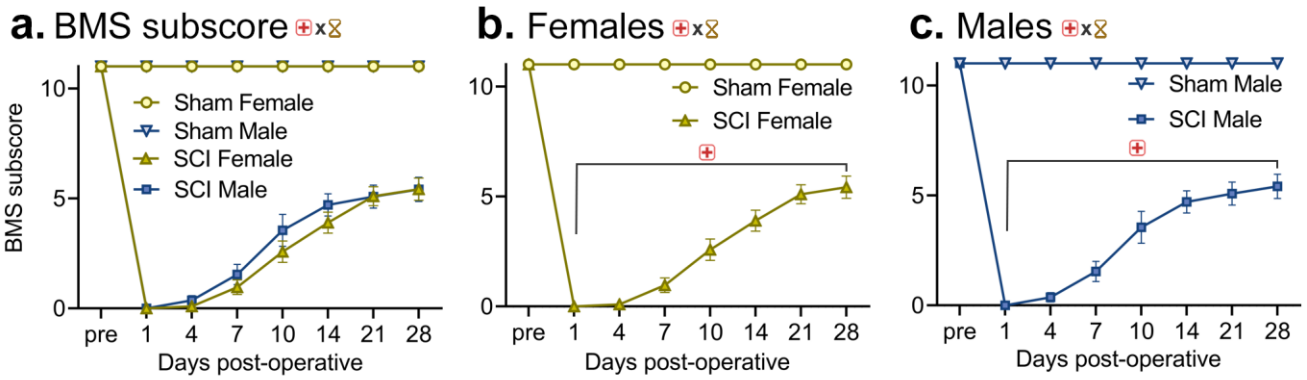


**Supplemental Figure 2.** T9 thoracic contusion SCI elicits similar locomotor deficits in female and male mice. Locomotor recovery after SCI or sham surgery in both sexes (**a**), females only (**b**), and males only (**c**). BMS subscores reveal similar recovery curves after SCI between sexes, although females showed slightly delayed recovery at acute times. Female-sham: n=39; male sham: n=48; female-SCI: n=33; male-SCI: n=24. + $p < 0.05$ between sham and SCI mice; hourglass symbol indicates $p < 0.05$ across timepoints; “x” symbol indicates significant interaction between respective groups. Symbols at the top of each panel indicate main effects. ANOVA (a: three-way; b,c: two-way repeated measures) with Holm-Sidak *post-hoc* test.
